# Deep Learning Based Tumor Type Classification Using Gene Expression Data

**DOI:** 10.1101/364323

**Authors:** Boyu Lyu, Anamul Haque

**Affiliations:** Virginia Tech Blacksburg, Virginia

**Keywords:** Deep Learning, Tumor Type Classification, Pan-Cancer Atlas, Convolutional Neural Network

## Abstract

Differential analysis occupies the most significant portion of the standard practices of RNA-Seq analysis. However, the conventional method is matching the tumor samples to the normal samples, which are both from the same tumor type. The output using such method would fail in differentiating tumor types because it lacks the knowledge from other tumor types. Pan-Cancer Atlas provides us with abundant information on 33 prevalent tumor types which could be used as prior knowledge to generate tumor-specific biomarkers. In this paper, we embedded the high dimensional RNA-Seq data into 2-D images and used a convolutional neural network to make classification of the 33 tumor types. The final accuracy we got was 95.59%, higher than another paper applying GA/KNN method on the same dataset. Based on the idea of Guided Grad Cam, as to each class, we generated significance heat-map for all the genes. By doing functional analysis on the genes with high intensities in the heat-maps, we validated that these top genes are related to tumor-specific pathways, and some of them have already been used as biomarkers, which proved the effectiveness of our method. As far as we know, we are the first to apply convolutional neural network on Pan-Cancer Atlas for classification, and we are also the first to match the significance of classification with the importance of genes. Our experiment results show that our method has a good performance and could also apply in other genomics data.

## 1 INTRODUCTION

The invention of Next Generation Sequencing methods has mostly boosted the analysis of human genomics due to the improvement in the efficiency and accuracy. To better understand the cause of various tumors, a large volume of tumor tissues has been sequenced and managed by The Cancer Genome Atlas (TCGA). With these samples, TCGA further analyzed over 11,000 tumors from 33 most prevalent forms of cancer, which fostered the accomplishment of Pan-Cancer Atlas. Where, as to each tumor sample, we could access its RNA-Seq expression data. These data are beneficial when trying to identify potential biomarkers for each tumor.

As to biomarkers, most analyses tried to find the differentially expressed genes. However, they didnt consider expressions of other tumors. Also, during the study, models are constructed to mimic the expression data. However, such models are very data specific, which would fail in dealing with data from other samples or other tumor types. So, it is highly needed to build a method which can include knowledge from multiple tumor types into the analysis.

On the other hand, tumor type classification can contribute to a faster and more accurate diagnosis. However, research on tumor type classification is currently rare, which is partially due to the difficulty in dealing with high dimensionality of the genomic dataset. In Pan-Cancer Atlas, the normalized mRNA-seq gene expression data contains information from more than 20k genes. Within that many genes, a lot of genes are actually of no specific effect on the tumor. These genes are weak features. Using generic machine learning methods such as KNN wont work due to the curse of dimension. Even though classification of tumor type is still in its beginning, deep learning has been vastly used in image classification/ recognition, some famous architectures like Resnet [9] and inception[17] have excellent performance. In the Imagenet 2017 challenge, the winner obtained a top-5 error rate of 2.251%[10] using a convolutional neural network. Besides, to understand how deep neural network works, in computer vision field, various visualization methods have been invented such as deep Taylor decomposition, layer-wise decomposition, and Grad-Cam[16]. They generate heat-maps in the input image to indicate the contribution of each pixel to the classification. Since the gene expression data contains more than 10k samples in the atlas, it is promising to have good accuracy with the deep neural network. At the same time, assuming that the importance of genes equals to the significance of genes in classification, we could borrow the interpretation methods of the deep neural network to discover top genes for each tumor. In this way, each gene will be assigned a confidence score, and genes with high scores are considered to be the top genes or potential biomarkers since their existence affect the classification most.

Based on the deep neural network and the visualization methods, we first filter out the genes with small variance across all the samples. Then, we embedded the high-dimension expression data (10381×1) into a 2-D image (102×102) to fit for the convolutional layers. Next, we constructed a three-layer convolution neural network and used 10-fold cross-validation to test the performance. With the trained neural network, we followed the idea of Guided Grad-Cam[16] and generated heat-maps for all the classes showing prominent pixels (genes). By functional analysis, we validated that the top genes selected in this way are biologically meaningful for corresponding tumors, which proves the effectiveness of our work.

The remaining of this paper is organized as follows. In section 2, we reviewed the related work of tumor type classification and visualization of the deep neural network. In section 3, we described the data we used, and the process of we applied for tumor type classification. Also, the steps to generate heat map for each sample are also explained. Finally, in section 4, The accuracy of the tumor type classification is discussed and compared with the result of GA/KNN method in paper[11]. Also, top genes are picked from the heat-maps and validated by functional analysis.

## 2 RELATED WORK

Binary classification (identification) of tumors has been found in some papers, but only paper [11] researched multiple tumor type classification using gene expression data. The authors applied GA/KNN method to iteratively generate the subset of the genes (features) and then use KNN method to test the accuracy. After hundreds of iterations, they obtained the feature set corresponding to the best accuracy. In this way, this method achieved an accuracy of 90% across 31 tumor types, and also generated a set of top genes for all the tumor type. GA/KNN method is a straightforward method which could obtain an optimal feature set in the end, but it requires running a lot of iterations and also fixes the size of feature set at first. However, using one single feature set to make classification overlooked the heterogeneity among different tumor types. Also, top genes for various tumor types could be varied a lot.

Deep learning method was also used to identify top genes and make cancer identification of one single type. In paper [5], the author used a stacked auto-encoder first to extract high-level features from the expression values and then input these features into a single layer ANN network to decide whether the sample is a tumor or not. The accuracy using such method reached 94%. However, as to multiple classes, this method will lead to a very complicated structure, since this is not an end-to-end method. To identify the top genes of BRCA, weight matrices of each layer in the auto-encoder are multiplied to obtain the estimated weights for each gene at the input layer. By fitting with the normal distribution, they selected the top genes. This idea is similar to our view of using visualization methods. The difference is that they matched the importance of genes to the high-level features, whereas we matched the importance of genes to their contribution to the classification.

Deep Taylor decomposition and layer-wise relevance propagation (LRP) [1, 2] are designed to interpret the deep neural network by backpropagation. Where one conservative way to perform LRP is to redistribute the input of each neuron back to all of its predecessors equally. And layer by layer, when the input layer is reached, the decomposition is done. However, LRP methods require a relevance conservation property between layers, which puts some constraints on the neural network. While another technique, guided Grad-Cam[16], a combination of Guided back-propagation and Grad-Cam, can be applied on any neural network without any modification to the original architecture. In this paper, we are going to use this method to trace the significant pixels (genes) in the input, so that we could extract substantial genes. The localization map of Grad Cam is generated by first calculating the activation map in the forward propagation and the gradient map in the backward propagation. Since the deeper the convolutional layer, the higher level feature it contains. And after sending into the fully connected layer, all the spatial information will be lost. So, Grad-Cam constructs the localization map of the last convolutional layer and resizes it to the input size. However, as stated by the [5], the heat map shows the importance of each class, but the resolution is low, due to the resizing process. To solve this problem, a pixel-space gradient visualization method, guided back propagation, was used to refine the heat map. The output of guided backpropagation is the gradient of each pixel in the input image, which can be obtained through the backpropagation. Then the final output of this Guided Gram-Cam can be obtained by multiplying the results of guided backpropagation and Grad-Cam.

## 3 DATA AND METHOD

### 3.1 Classification

The workflow of our method is shown in figure1. (1) Preprocessing of the input data. (2) Make tumor type classification using a convolutional neural network. (3) Generate the heat map for each class and pick the genes corresponding to top intensities in the heat-maps. (4) Validate the pathways of selected genes.

**Figure 1:**
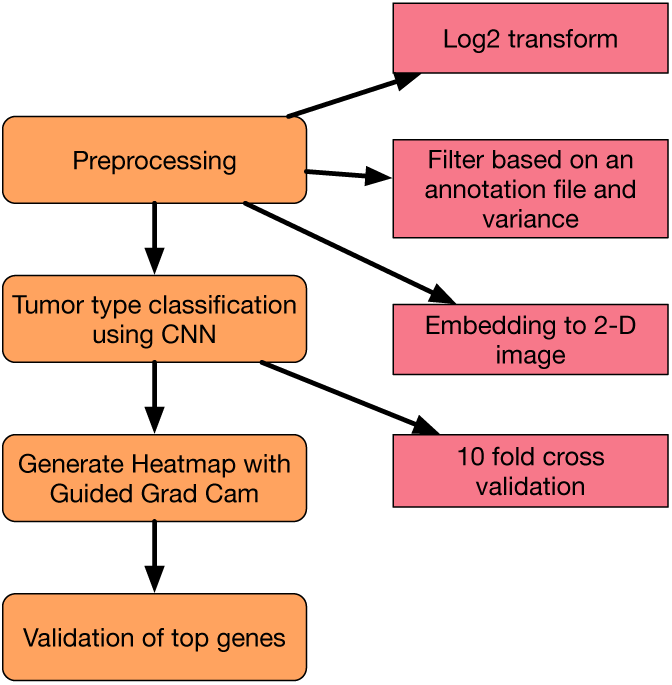
The workflow of our method.

### 3.2 Data

We used the normalized-level3 RNA-Seq gene expression data of 33 tumor types in Pan-Cancer Atlas. A detailed list of all 33 tumor types and corresponding number of samples is shown in table 3. The data contains 10267 tumor samples with respect to 20531 genes.

### 3.3 Preprocessing

The data contains the normalized read count for each gene, but the range of the values is enormous, and also some values are smaller than 1. So we first apply a transform using *y* = log_2_(*x* + 1), and set all the values lower than 1 to be 0 since these values are very likely to be noise. Besides, we matched the genes with an annotation file (updated on 04/03/2018) downloaded from NCBI, around 1000 genes were not found in the annotation file so that we removed these genes and corresponding expression level. Then we used a variance threshold of **1.19** to filter out the genes of which expression levels remain almost unchanged, and the number of genes is further reduced to 10381.

To input the data to the convolutional neural network, we embedded the data into 2-D images. First genes are ordered based on the chromosome number because adjacent genes are more likely to interact with each other. Then the data is reshaped from a 10381 × 1 array into a 102 × 102 image by adding some zeros at the last line of the image. Then all the images are normalized to make sure that the range is [0,255]. The generated images of different classes are shown in figure 2.

**Figure 2:**
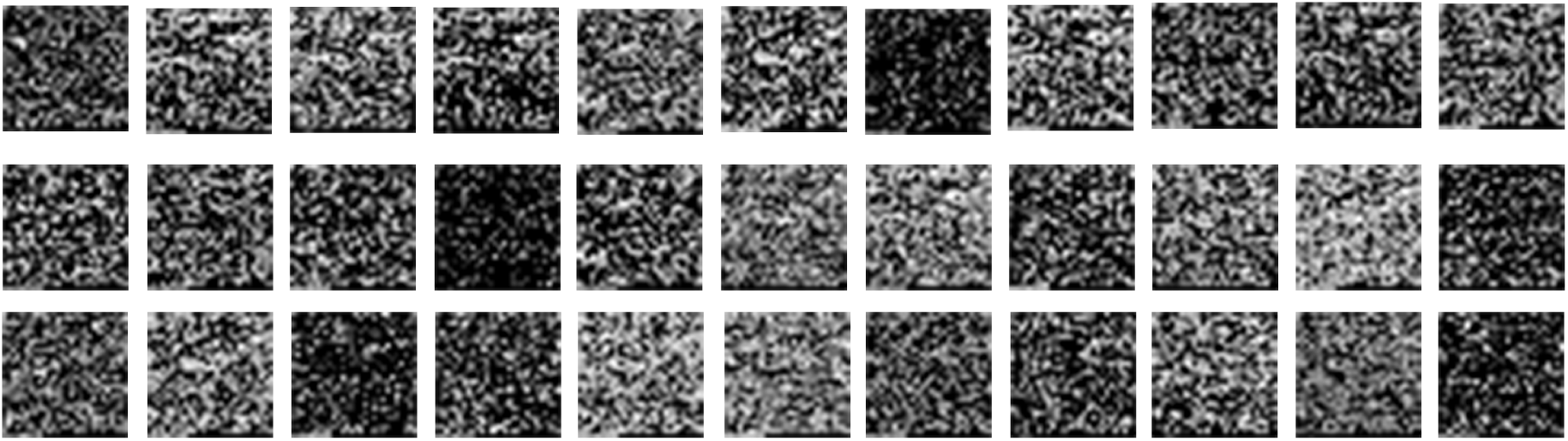
An example of embedded 2-D images. From top left to the bottom right are the images generated from class 1 to class 33.

### 3.4 Classification

We used a convolutional neural network consisting of three convolutional layers, and three fully connected layer, which is shown in figure 5. The first convolutional layer ‘conv1’ contains 64 different filters, while the second and the third convolutional layers contain 128 and 256 filters respectively. Max-pooling layer and batch-normalization layer are followed immediately after each convolutional layer. A drop-out layer is added before entering into fully connected layer, the drop-out rate is 25%. The size of the three fully connected layers are 36864, 1024, 512 separately. We chose Cross Entropy as loss function and Adam optimizer to update the weights. We used 10-fold cross validation to train the convolutional neural network and to test the performance.

### 3.5 Heat-map

Based on the idea of Guided Grad-Cam, we let the training data set go through the trained neural network. We recorded the activation maps in the forward pass and the gradient maps in the backpropagation, then we could obtain the heat map for each sample using Guided Grad-Cam. And by averaging all the heat-maps from the same class and after normalization, we could obtain the class specific heat-map. As to each pixel in the heat-map, a higher intensity represents a higher contribution to the final classification, which indicates a higher importance of its corresponding gene.

In fact, this heat-map generating process could be implanted into the training of the convolutional neural network, because it will not affect the training and testing process. But the computation is less using the above two-step method, because Grad-Cam only requires all the samples to run through the neural network once.

### 3.6 Validation

Top genes are selected based on the intensity rankings in the heat-maps. We applied functional analysis on these top genes to further prove that the genes are tumor specific and are potential biomarkers. In the first stage, we chose top 400 genes of each tumor type to do pathway analysis, trying to find out if significantly enriched pathways are related to the corresponding tumor. In the second stage, we studied the top 5 genes for the tumor types presenting few results in the first stage, to find their relations to the tumor.

## 4 EXPERIMENT RESULT

### 4.1 Classification

Using 10-fold cross validation, we calculated the average accuracy, average accuracy of each class, average precision score *P* and average recall score *R*, and average F1 score *F*1. Where,

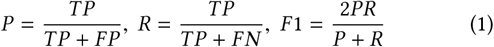

The scores are summarized into table 1

**Table 1:** Performance of our method.

| method | accuracy | precision | recall | f1-score |
| --- | --- | --- | --- | --- |
| CNN | 95.59% | 95.54% | 95.59% | 95.43% |

In addition, we have also generated a confusion matrix as shown in figure3 We can see most classes are classified correctly, but there are two pairs of misclassification. (1) READ samples are mostly misclassified into COAD which might be due to the close spatial relationship between the two tumor types. (2) Some CHOL samples are misclassified into LIHC due to the small number of CHOL samples. But using CNN do show an improvement in dealing with READ samples compared to the reference[11]. A comparison of accuracy as to each class is shown in table 3. From the table we can see that our accuracies in dealing with classes READ and UCS have been improved a lot compared with the reference paper.

**Figure 3:**
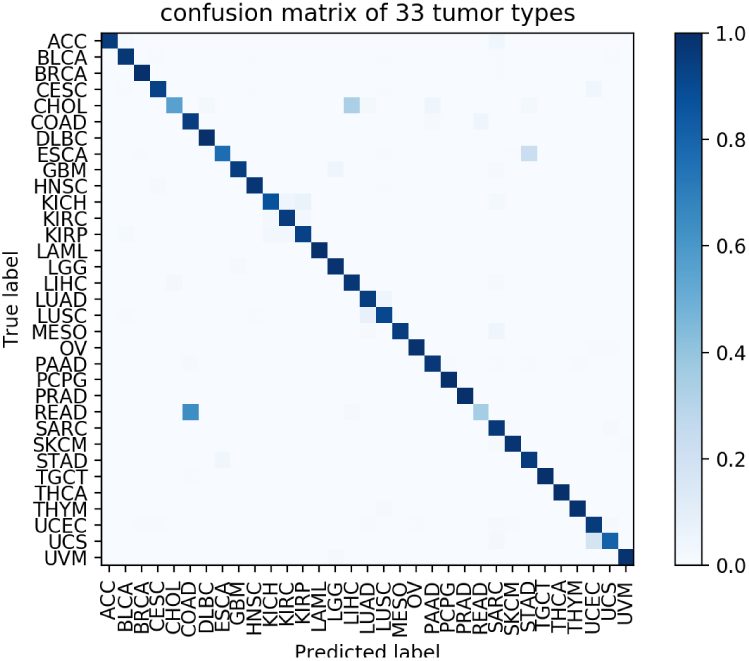
The confusion matrix generated using our method.

### 4.2 Generated Heat-map

The heat-maps generated for each class showed a similarity across ten folds, and displayed a distinct pattern when comparing in-between classes. Some examples are shown in figure 6. In the example, each row represents heat-maps of one tumor type, and columns represent heat-maps from different folds (from left to right are fold 1 to fold 10). Circled regions show the similar pattern in all the folds. Even though there are some difference among different folds, there do exist a clear pattern.

### 4.3 Validation of Top Genes

In the heat-maps, we found that the intensities of top 100 genes decreased sharply, while the intensities decreased very smoothly starting from around 400th genes. When the intensities are very small, the decrease rate became high again. Such change can be viewed in fig 4. Assuming that the larger intensity implies more significance, since the slopes of the intensity change in the first 400 genes are larger than the following several thousands genes, we chose the top 400 genes as query genes. Besides, comparing to the total number of genes (10381), our choice of 400 is still a very small number, which is consistent with the fact that the number of biomarkers should be small. The KEGG pathway analysis results for top 400 genes of each tumor are obtained using the David website^1^. Next, we did a literature review on the results trying to find out the relations between these pathways and tumor types. The related pathways with P value smaller than 10^−3^ are shown in table4. Where the bold ones are related to corresponding tumor types. The top 400 genes of 24 tumor types were found to have significant enrichments of pathways. In which, 16 tumor types have at least one related pathway given the top 400 genes. The related genes in these pathways can then be viewed as tumor specific biomarkers.

**Figure 4:**
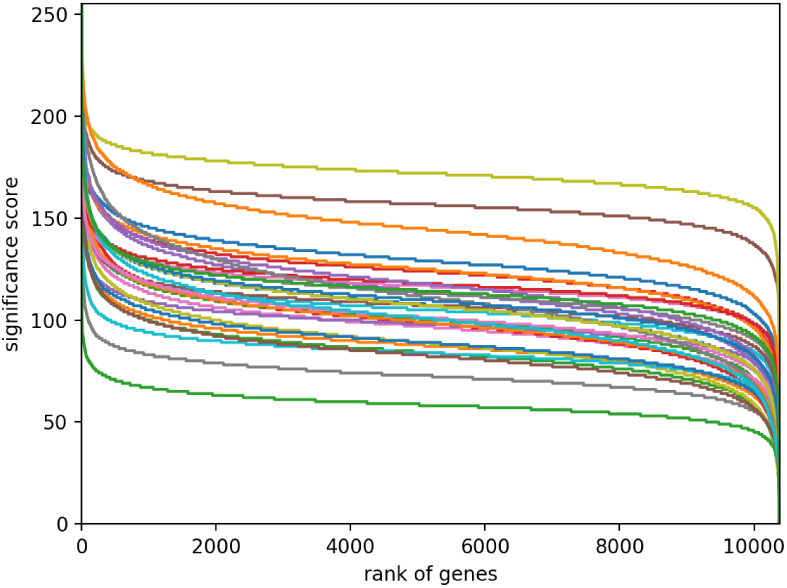
The changes of intensities in heat-maps for each class. It can be seen that different classes shared the same pattern of change in intensities.

**Figure 5:**
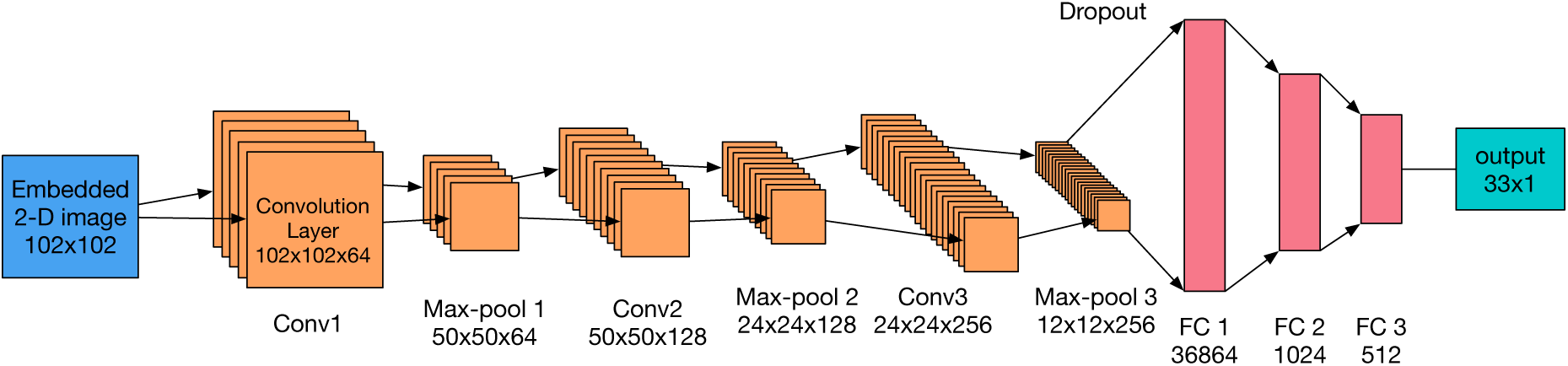
The architecture of our convolutional neural network.

**Figure 6:**
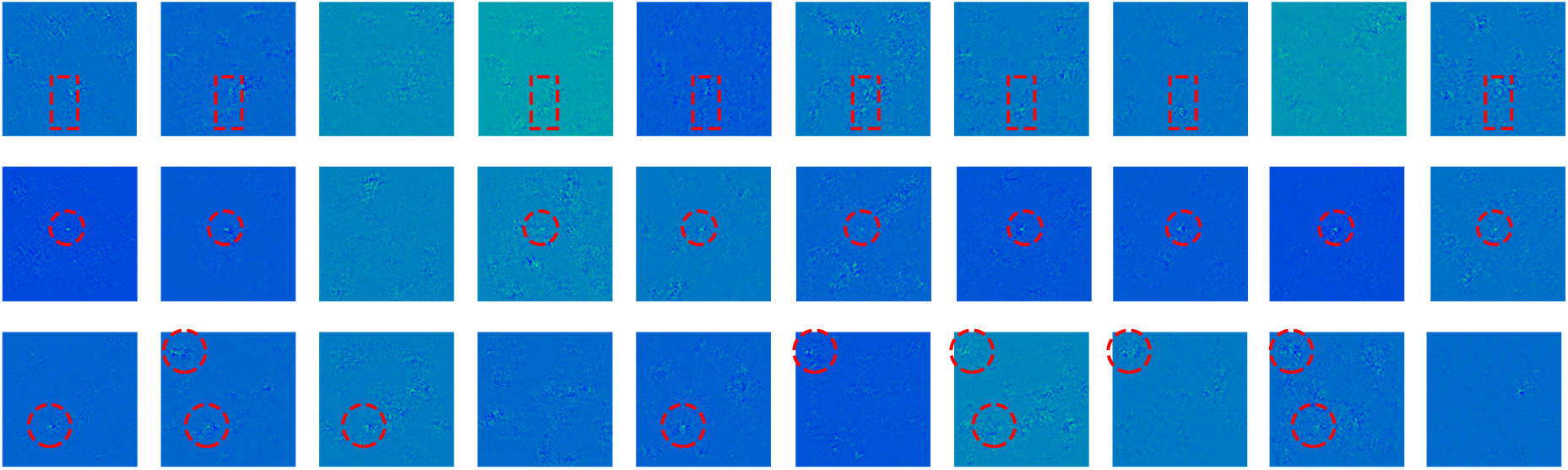
Some heat map examples. Each column represents the result from one fold. The first row are the heat-map of tumor type BLCA, the second row belongs to PRAD and the third row belongs to LGG.

While as to the other 8 tumor types, even though no direct related pathways are found, two significant enriched concurrent pathways are discovered, which are hsa04512: ECM-receptor interaction and hsa04510: focal adhesion. However, the related genes are not the same as to different tumor types. For example, in BRCA, ECM-receptor interaction pathway related genes are SDC1, COL4A1, COL3A1, COL6A3, COL1A2, ITGA11, COL6A2, COL6A1, COL1A1, COL5A2 and FN1. Whereas in ESCA, related genes are VWF, ITGA6, LAMC3, ITGAV, TNC, ITGB6, ITGB4, ITGA3, COL1A1, COL5A2 and COL4A6. Similarly, in terms of focal adhesion pathway, related top genes in SKCM are CAV1, TNC, COL3A1, PIK3CD, ITGA3, ITGB3, COL5A3, CCND1, LAMB2, FYN, COMP, COL6A3, COL6A2, COL6A1, COL1A1, LAMB1, PARVB, SPP1, FN1 and SHC4. Whereas related genes in THCA are COL4A4, COL4A3, CAV2, CAV1, PGF, BCAR1, MET, IGF1, ITGA3, LAMA5, COL1A2, LAMC2, COL1A1 and FN1. It can be seen that these sets are not quite the same in different tumor types, which shows that they can be potential biomarkers.

In the other 9 tumor types (CHOL, COAD, READ, GBM, KICH, LGG, LUSC, OV and UVM), no significant enriched pathways were found. The samples for CHOL and KICH are limited, so we omitted these two tumor types. Besides, the classification accuracy of class READ is very low, so we also omitted it. With respect to the 6 remaining tumor types, we reduced the query size from top 400 to top 5. The query genes are shown in table 2. We looked into their information on the GeneCards website^2^. As to COAD, its top1 gene LGALS4 (Galectin 4) is a Protein Coding gene. The expression of this gene is restricted to small intestine, colon, and rectum, and it is under-expressed in colorectal cancer. As to GBM (Glioblastoma multiforme), a cancer in brain region, its top1 gene GFAP (Glial Fibrillary Acidic Protein) is a Protein Coding gene. This gene encodes one of the major intermediate filament proteins of mature astrocytes. It is used as a marker to distinguish astrocytes from other glial cells during development, which is also in the brain region. LGG (Brain Lower Grade Glioma) is also a tumor in the brain, its top1-4 genes are all pseudo-genes, while the top5 gene GFAP is the gene related to brain. As to LUSC (Lung squamous cell carcinoma), its top1 gene SFTPA2 has been implicated in many lung diseases [13]. As to OV (Ovarian cancer), in paper[7], its top1 gene MUC16 (CA125) was said to be the **only** reliable diagnostic marker for ovarian cancer. As to UVM (Uveal Melanoma), its top1 gene CD44 were tested to be strongly expressed in several cell lines of human uveal melanoma[4]. All the above results have shown the top genes have very close relations to the corresponding tumor types, which could be viewed as potential biomarkers.

**Table 2:** Top5 genes for the 6 remaining tumor types.

| Rank | COAD | GBM | LGG |
| --- | --- | --- | --- |
| 1 | LGALS4 | GFAP | HSPB1P1 |
| 2 | FCGBP | CBR1 | HNRNPA1P33 |
| 3 | FTL | LOC613037 | EEF1A1P9 |
| 4 | HOXC6 | TIMP2 | FTHL3 |
| 5 | IGFBP2 | IGFBP5 | GFAP |
| Rank | LUSC | OV | UVM |
| 1 | SFTPA2 | MUC16 | CD44 |
| 2 | KRT6A | KLK6 | LGALS3BP |
| 3 | SFTPA1 | KLK7 | SERPINF1 |
| 4 | SFTPB | KLK8 | GAPDHS |
| 5 | KRT6B | KCNK15 | MGST3 |

**Table 3:**
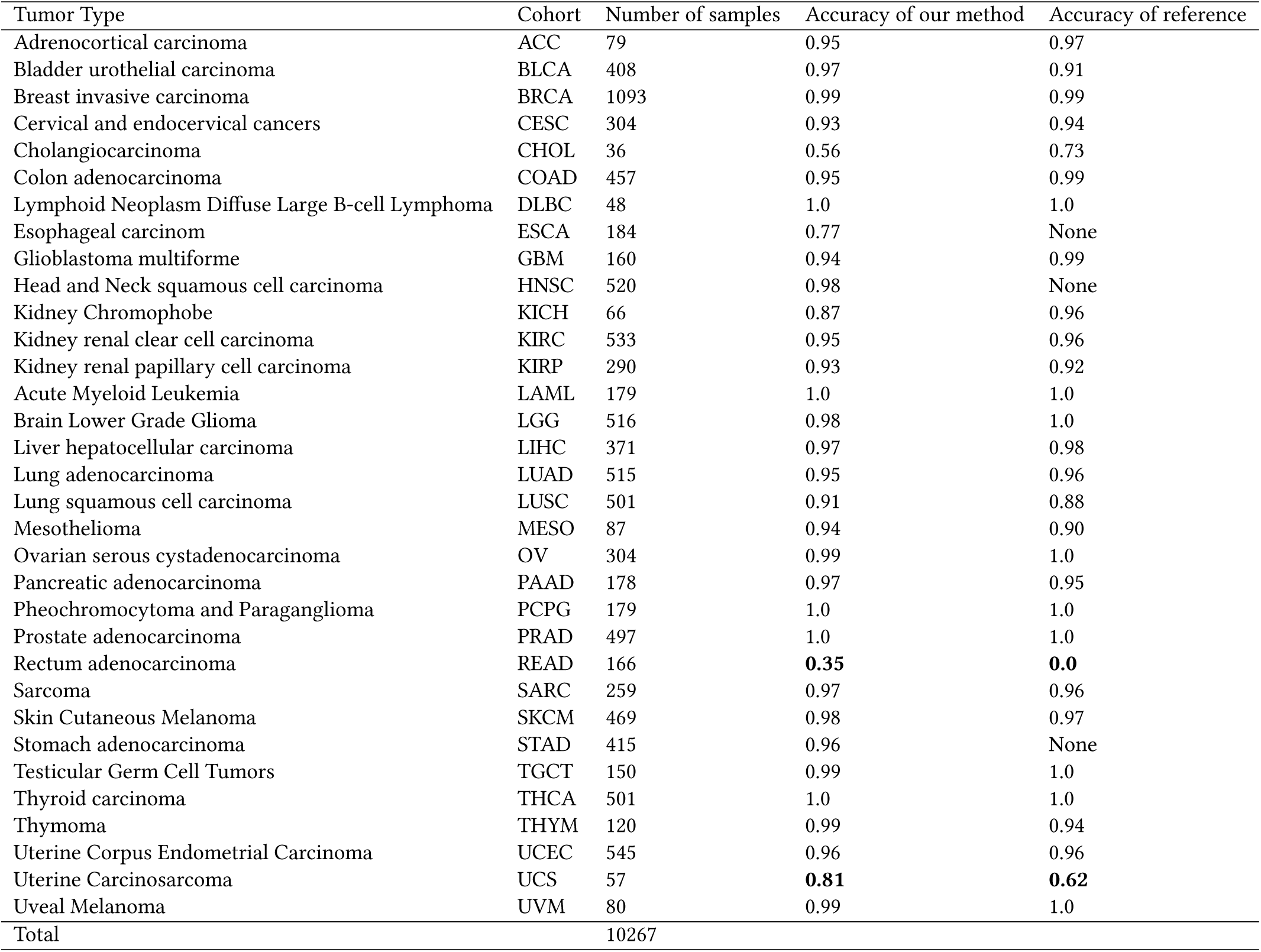
Tumor types and number of RNA-Seq samples.

**Table 4:** Pathway analysis results on top 400 genes of each tumor type (*P* < 10^−3^)

| Tumor name | Related Pathway |  | P value | Tumor name | Related Pathway |  | P value |
| --- | --- | --- | --- | --- | --- | --- | --- |
|  | ID | Name |  |  | ID | Name |  |
| ACC | **hsa04925** | **Aldosterone synthesis and secretion** | 3.13E-06 | LAML | **hsa04672** | **Intestinal immune network for IgA production** | 1.27E-04 |
|  | hsa04913 | Ovarian steroidogenesis | 1.83E-05 |  | **hsa05140** | **Leishmaniasis[20]** | 3.11E-04 |
|  | hsa01100 | Metabolic pathways | 4.69E-04 | LIHC | hsa04610 | Complement and coagulation cascades | 6.85E-15 |
| BLCA | **hsa00140** | **Steroid hormone biosynthesis[12]** | 5.63E-07 |  | hsa05150 | Staphylococcus aureus infection | 3.25E-05 |
|  | hsa04510 | Focal adhesion | 3.69E-06 |  | hsa00830 | Retinol metabolism | 7.71E-04 |
|  | hsa04512 | ECM-receptor interaction | 3.71E-06 | LUAD | hsa00512 | Mucin type O-Glycan biosynthesis | 4.57E-04 |
|  | hsa04974 | Protein digestion and absorption | 4.16E-06 | MESO | hsa04512 | ECM-receptor interaction | 7.29E-10 |
|  | hsa00980 | Metabolism of xenobiotics by cytochrome P450 | 2.51E-04 |  | hsa04610 | Complement and coagulation cascades | 3.37E-09 |
| BRCA | hsa04512 | ECM-receptor interaction | 1.30E-05 |  | **hsa05150** | **Staphylococcus aureus infection** | 2.42E-08 |
|  | hsa04510 | Focal adhesion | 1.05E-04 |  | hsa04974 | Protein digestion and absorption | 5.75E-07 |
| CESC | **hsa05205** | **Proteoglycans in cancer[14]** | 3.10E-04 |  | hsa04510 | Focal adhesion | 8.13E-07 |
| DLBC | **hsa05330** | **Allograft rejection[8]** | 3.91E-17 |  | **hsa04151** | **PI3K-Akt signaling pathway[15]** | 9.60E-05 |
|  | **hsa05332** | **Graft-versus-host disease[3]** | 1.15E-16 |  | hsa04145 | Phagosome | 1.73E-04 |
|  | hsa04940 | Type I diabetes mellitus | 5.47E-16 |  | **hsa04611** | **Platelet activation[15]** | 7.07E-04 |
|  | hsa05320 | Autoimmune thyroid disease | 3.78E-14 | PAAD | hsa04974 | Protein digestion and absorption | 3.48E-09 |
|  | hsa05150 | Staphylococcus aureus infection | 7.73E-14 |  | hsa04950 | Maturity onset diabetes of the young | 8.01E-07 |
|  | hsa04672 | Intestinal immune network for IgA production | 1.06E-13 |  | hsa04512 | ECM-receptor interaction | 1.38E-05 |
|  | hsa05416 | Viral myocarditis | 2.12E-13 | PCPG | hsa04721 | Synaptic vesicle cycle | 4.53E-05 |
|  | hsa04612 | Antigen processing and presentation | 2.70E-13 | PRAD | **hsa04270** | **Vascular smooth muscle contraction** | 2.69E-04 |
|  | hsa04145 | Phagosome | 1.31E-12 | SARC | hsa04512 | ECM-receptor interaction | 5.42E-11 |
|  | hsa05310 | Asthma 13 | 1.53E-11 |  | hsa04974 | Protein digestion and absorption | 7.52E-10 |
|  | hsa05321 | Inflammatory bowel disease (IBD) | 2.23E-11 |  | hsa04510 | Focal adhesion | 1.32E-09 |
|  | hsa05166 | HTLV-I infection | 4.38E-11 |  | **hsa05146** | **Amoebiasis** | 2.89E-05 |
|  | hsa05323 | Rheumatoid arthritis | 4.57E-11 |  | hsa04151 | PI3K-Akt signaling pathway | 2.55E-04 |
|  | hsa05140 | Leishmaniasis | 1.22E-10 | SKCM | hsa04512 | ECM-receptor interaction | 3.08E-08 |
|  | hsa05168 | Herpes simplex infection | 2.50E-09 |  | hsa04510 | Focal adhesion | 6.44E-08 |
|  | **hsa04514** | **Cell adhesion molecules (CAMs)** | 2.36E-08 |  | hsa04974 | Protein digestion and absorption | 3.02E-07 |
|  | **hsa04064** | **NF-kappa B signaling pathway** | 1.75E-07 |  | hsa05222 | Small cell lung cancer | 7.66E-05 |
|  | hsa05152 | Tuberculosis | 1.77E-07 |  | hsa05200 | Pathways in cancer | 8.55E-05 |
|  | **hsa05340** | **Primary immunodeficiency** | 3.63E-07 |  | hsa04151 | PI3K-Akt signaling pathway | 1.29E-04 |
|  | hsa04060 | Cytokine-cytokine receptor interaction | 7.92E-07 | STAD | **hsa04950** | **Maturity onset diabetes of the young[19]** | 9.03E-04 |
|  | hsa05145 | Toxoplasmosis | 7.60E-06 | TGCT | hsa04974 | Protein digestion and absorption | 4.96E-10 |
|  | **hsa05322** | **Systemic lupus erythematosus** | 3.30E-05 |  | hsa04512 | ECM-receptor interaction | 1.74E-05 |
|  | **hsa04062** | **Chemokine signaling pathway** | 9.90E-05 | THCA | hsa04512 | ECM-receptor interaction | 1.15E-05 |
|  | hsa05164 | Influenza A | 1.60E-04 |  | hsa04510 | Focal adhesion | 3.40E-04 |
|  | **hsa04662** | **B cell receptor signaling pathway** | 1.62E-04 |  | hsa04918 | Thyroid hormone synthesis | 5.91E-04 |
|  | hsa04640 | Hematopoietic cell lineage | 1.69E-04 | THYM | hsa04640 | Hematopoietic cell lineage | 6.43E-09 |
|  | **hsa04666** | **Fc gamma R-mediated phagocytosis** | 7.20E-04 |  | **hsa05340** | **Primary immunodeficiency[18]** | 2.67E-04 |
| ESCA | hsa04512 | ECM-receptor interaction | 2.40E-06 |  | hsa05162 | Measles | 9.46E-04 |
|  | hsa04510 | Focal adhesion | 1.28E-05 | UCEC | **hsa04530** | **Tight junction** | 5.60E-05 |
|  | hsa05412 | Arrhythmogenic right ventricular cardiomyopathy (ARVC) | 2.97E-05 | UCS | hsa04512 | ECM-receptor interaction | 1.02E-04 |
|  | hsa04974 | Protein digestion and absorption | 1.40E-04 |  | hsa04974 | Protein digestion and absorption | 1.12E-04 |
| HNSC | **hsa05414** | **Dilated cardiomyopathy[6]** | 2.01E-06 |  | hsa04510 | Focal adhesion | 5.05E-04 |
|  | hsa04512 | ECM-receptor interaction | 2.78E-06 | KIRP | hsa04512 | ECM-receptor interaction | 5.93E-05 |
|  | hsa05410 | Hypertrophic cardiomyopathy (HCM) | 8.68E-06 |  | **hsa00040** | **Pentose and glucuronate interconversions** | 2.43E-04 |
|  | hsa05412 | Arrhythmogenic right ventricular cardiomyopathy (ARVC) | 2.47E-04 |  | hsa00053 | Ascorbate and aldarate metabolism | 4.92E-04 |
|  | hsa04261 | Adrenergic signaling in cardiomyocytes | 2.54E-04 |  | **hsa00140** | **Steroid hormone biosynthesis** | 5.79E-04 |
|  | hsa04260 | Cardiac muscle contraction | 3.47E-04 | KIRC | **hsa04610** | **Complement and coagulation cascades** | 1.21E-06 |

## 5 DISCUSSION

Genomic tests are becoming popular nowadays. With saliva or blood, these tests can tell out the estimated possibilities of different tumors for each individual. Such tests mostly rely on biomarkers of various tumors. However, gene expression biomarkers generated by differential analysis are not guaranteed to be tumor specific, since several tumors might share the same biomarkers. To avoid this problem, we designed a method using the knowledge from multiple tumor types and found the genes which can be used to differentiate between different tumor types. Validation results prove that these top genes are possible to be biomarker due to their relations to the corresponding tumor type. Further examinations can be made biologically.

In this paper, we used a convolutional neural network to make classification on the genomics data. Research on the computer vision is developing very fast. A lot of methods were designed to solve problems using the deep neural network. However, in the genomics community, the research based on deep learning is still in its beginning. One issue is that genomics data usually are high dimensional, while most deep learning architectures are for 2-D images. While, we have shown that, by just naively placing the genomics data (genes) onto each pixel of the 2-D image based on the order of chromosome number, the performance was excellent except for several tumor types. So that, it is possible that many more deep learning methods could be applied to the genomics.

Looking into the misclassified samples, we think one possible issue is the imbalanced dataset. Some tumor type has over 1000 samples while some only has 30 samples. Imbalanced dataset might cause a big problem in deep learning, so for a better result, this problem can be mitigated using oversampling methods such as SMOTE.

## 6 CONCLUSION

In this paper, we designed a new method to discover potential biomarkers for each tumor type. We matched the importance of genes to the contribution to the correct classification. Based on the Pan-Cancer Atlas, we used a convolutional neural network to make the classification and used a visualization method of neural network to discover top genes from the input. The accuracy of our method shows improvement compared with previous work on tumor type classification. And we examined the top genes of each tumor type, found their relations to the corresponding tumor types, which proved that the top genes are potential biomarkers.

## Footnotes

1 David:https://david.ncifcrf.gov/home.jsp

2 GeneCards: www.genecards.org

